# Single-cell Analysis of ACE2 Expression in Human Kidneys and Bladders Reveals a Potential Route of 2019-nCoV Infection

**DOI:** 10.1101/2020.02.08.939892

**Authors:** Wei Lin, Longfei Hu, Yan Zhang, Joshua D. Ooi, Ting Meng, Peng Jin, Xiang Ding, Longkai Peng, Lei Song, Zhou Xiao, Xiang Ao, Xiangcheng Xiao, Qiaoling Zhou, Ping Xiao, Jue Fan, Yong Zhong

**Author notes:** These authors contribute equally to this work (Wei Lin, Longfei Hu). Correspondence to: Jue Fan, Yong Zhong.

## Abstract

Since December 2019, a novel coronavirus named 2019 coronavirus (2019-nCoV) has emerged in Wuhan of China and spread to several countries worldwide within just one month. Apart from fever and respiratory complications, acute kidney injury has been observed in some patients with 2019-nCoV. In a short period of time, angiotensin converting enzyme II (ACE2), have been proposed to serve as the receptor for the entry of 2019-nCoV, which is the same for severe acute respiratory syndrome coronavirus (SARS). To investigate the possible cause of kidney damage in 2019-nCoV patients, we used both published kidney and bladder cell atlas data and an independent unpublished kidney single cell RNA-Seq data generated in-house to evaluate ACE2 gene expressions in all cell types in healthy kidneys and bladders.

Our results showed the enriched expression of all subtypes of proximal tubule cells of kidney and low but detectable levels of expression in bladder epithelial cells. These results indicated the urinary system is a potential route for 2019-nCoV infection, along with the respiratory system and digestion system. Our findings suggested the kidney abnormalities of SARS and 2019-nCoV patients may be due to proximal tubule cells damage and subsequent systematic inflammatory response induced kidney injury. Beyond that, laboratory tests of viruses and related indicators in urine may be needed in some special patients of 2019-nCoV.

## Introduction

In late December 2019, a cluster of mystery pneumonia cases emerged in Wuhan, a city in the middle south of China. Deep sequencing analysis and etiological investigations then confirmed the pathogen was a type of a newly identified coronavirus which had been labelled as 2019 novel coronavirus (2019-nCoV). The rapid spread of 2019-nCoV caused the outbreak of pneumonia in China and also other countries, making it a severe threat to international public health security within a short period of time^1-3^.

Based on bioinformatics analyses, it has been shown that 2019-nCoV genome is 88% identical with two bat-derived severe acute respiratory syndrome (SARS)-like coronaviruses (bat-SL-CoVZC45 and bat-SL-CoVZXC21), 79% with SARS-CoV, and 50% with MERS-CoV^4^. Protein structural analyses revealed that 2019-nCoV had a similar receptor-binding domain to that of SARS-CoV, directly binding to angiotensin converting enzyme II (ACE2), strongly suggesting that 2019-nCoV uses ACE2 as its receptor^5-8^.

The most common symptoms of 2019-Cov infection are fever and cough in most patients, with symptoms of chest discomfort, progressive dyspnea or acute respiratory distress syndrome (ARDS) in severe patients. Acute kidney injury (AKI) has been reported in a small number of patients with confirmed 2019-nCoV infection. The laboratory findings of 99 patients infected with 2019-nCoV showed that 3 patients (3%) had different degrees of increased serum creatinine^9^. In another clinical observation study, 4 (10%) of all 41 patients, including 2 of 13 (15%) ICU and 2of 28 (7%) non-ICU patients, showed the elevated serum creatinine^10^. However, why and how the 2019-nCoV induced AKI remains largely unknown. In addition, whether 2019-nCoV can be transmitted through the urinary tract still remains a largely unanswered question. As ACE2 plays an important role in the 2019-nCoV infection, we aimed to identify the ACE2-expressing cell composition and proportion in normal human kidneys and bladders by single-cell transcriptomes in order to explore the possible infection routes of 2019-nCov and the roles of ACE2 in urinary tract system infection.

Single-cell RNA sequencing (scRNA-Seq) has been extensively applied in 2019-nCoV research, due to its capability to profile gene expressions for all cell types in multiple tissues unbiasedly at a high resolution. In addition to well-known alveolar type 2 cells in lung, expression of ACE2 has also been investigated in liver cholangiocytes, esophagus epithelial cells and absorptive enterocytes of ileum and colon by scRNA-Seq^11-13^. These results demonstrated the potential utility of scRNA-Seq to unmask the potential target cell types of 2019-nCoV. Since the urinary system infection and its potential aftermath could be essential to the patient care during and after the infection, here we used two scRNA-Seq transcriptome data in healthy kidneys and one dataset in heathy bladders to investigate the expression patterns of cell types in the urinary system.

## Methods

### Public dataset acquisition and processing

Gene expression matrices of scRNA-Seq data from normal kidneys of three healthy donors were downloaded from the Gene Expression Omnibus (GSE131685). We reproduced the downstream analysis using the code provided by the author in the original paper.

We obtained the scRNA-Seq data of healthy bladder tissues of three bladder cancer patients from the Gene Expression Omnibus (GSE108097). To be consistent with the kidney datasets, we applied Harmony^14^ to integrate samples and performed downstream analysis using Seurat V3. Clustering analysis was done by first reducing the gene expression matrix to the first 20 principal components and then using resolution 0.3 for the graph-based clustering. We annotated the cell types afresh using the canonical markers listed in the supplementary table of the original paper.

### Kidney sample processing

Kidney samples were obtained at a single site by wedge and needle biopsy of living donor kidneys after removal from the donor and before implantation in the recipient. Kidney biopsy samples were cleaned with sterile PBS after acquisition.

### Tissue dissociation

The fresh kidney tissue was immediately transferred into the GEXSCOPE Tissue Preservation Solution (Singleron Biotechnologies) at 2-8°C. The samples were digested in 2ml GEXSCOPE Tissue Dissociation Solution (Singleron Biotechnologies) at 37°C for 15min in a 15ml centrifuge tube with continuous agitation after washed with Hanks Balanced Salt Solution (HBSS) for three times and cut into approximately 1-2 mm pieces. Subsequently, cell debris and other impurities were filtered by a 40-micron sterile strainer (Corning). The cells were centrifuged at 1000 rpm and 4°C for 5 minutes. Cell pellets were resuspended into 1ml PBS (HyClone). To remove red blood cells, 2 mL GEXSCOPE Red Blood Cell Lysis Buffer (Singleron Biotechnologies) was added to the cell suspension and incubated at 25°C for 10 minutes. The mixture was then centrifuged at 1000 rpm for 5 min and the cell pellet resuspended in PBS. Cells were counted with TC20 automated cell counter (Bio-Rad).

### Library preparation and data analysis of scRNA-Seq

The single cell suspension was proceeded to the single cell library preparation and sequencing, as previously described^15^. To be consistent with the public data used above, we applied a similar downstream analysis workflow^14^. We modified the number of principal components to 20 and used resolution 0.6 to obtain comparable cell type clustering results with the public kidney data. Cell types were annotated based on the canonical markers in the literature ^14 16^.

## Results

### Expression patterns of ACE2 in kidney

By analyzing the public single-cell transcriptome dataset of normal human kidney cells from three donors^14^, we found ACE2 expression distributed across multiple cell types. Notably, ACE2 was mostly enriched in proximal tubule cells, including both convoluted tubule and straight tubule (Fig 1A-D). The other nephron subtypes, such as collecting duct and distal tubule as well as immune cells, all showed extremely low gene expressions.

**Figure 1.**
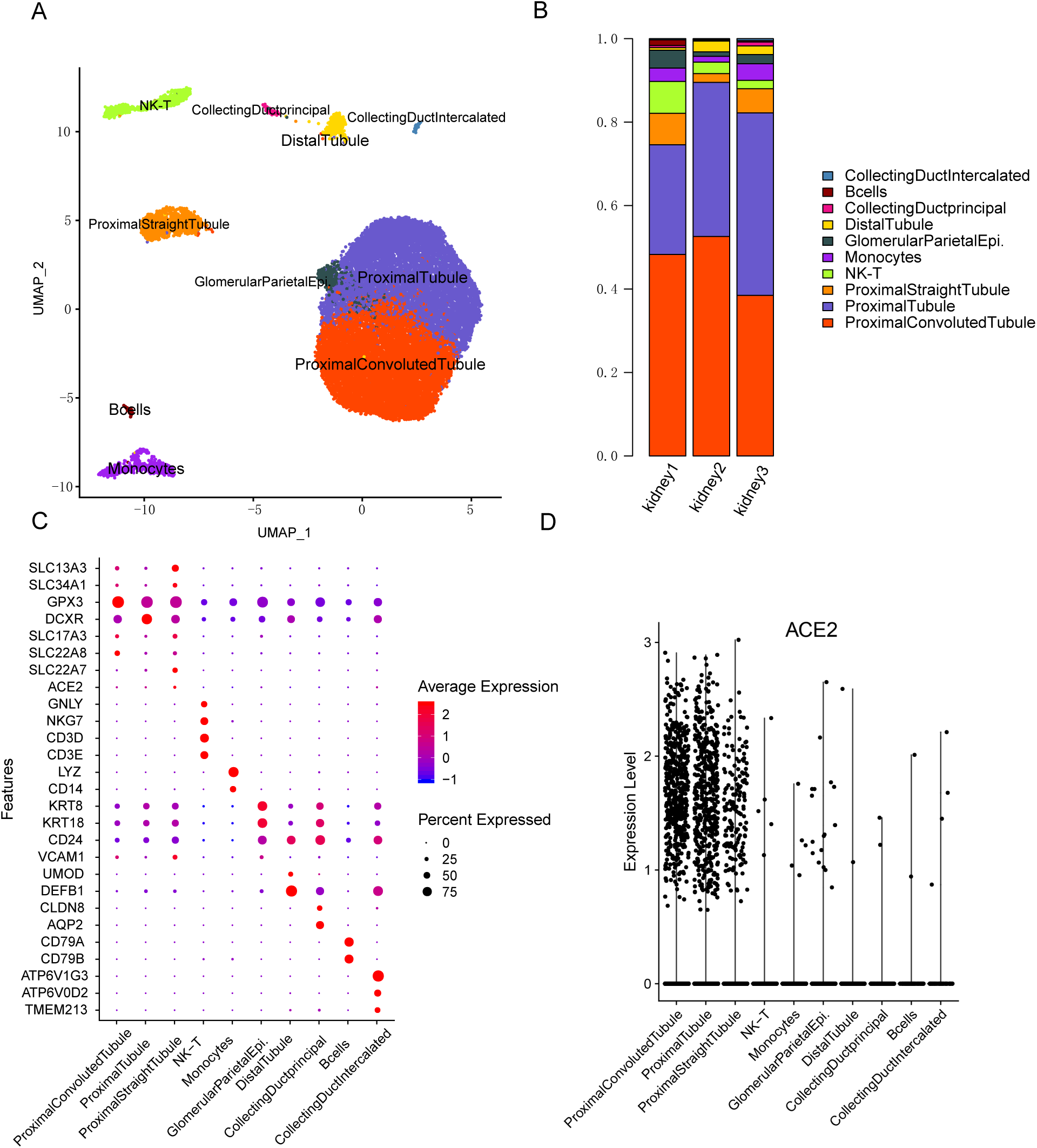
Single cell analysis of the public kidney data of three donors. A) UMAP plot revealing the cell type annotations of three samples. B) The cellular composition of three healthy kidney samples. C) Dotplot exhibition of canonical markers used for cell type classification. D) The gene expression patterns of ACE2 of all detected cell types in kidney.

To validate this result, we further analyzed normal kidney samples from 2 healthy donors. The 4736 cells were classified into 9 cell types, which were highly overlapped with the public data including 7 nephron specific subtypes based on the canonical marker genes (Fig 2A). The proximal tubule (PT) (CUBN, SLC13A3, SLC22A8) were classified into proximal convoluted tubule and proximal straight tubule based on different marker expression level^14 16^. Collecting duct principal cell (AQP2), collecting duct intercalated cells (ATP6V0D2) and glomerular parietal epithelial cells (KRT8, KRT18) were also annotated, in line with the previous result. Furthermore, we identified a cluster of loop of henle cells (UMOD, SLC12A1, CLDN16), which was absent in the public dataset. In addition, the stroma and immune cells, composed of a small number of endothelial cells (CD34, PECAM1) and monocytes (CD14, FCN1, VCAN), were found (Fig 2B). The representative marker genes for all subpopulations were plotted in Fig2C.

**Figure 2.**
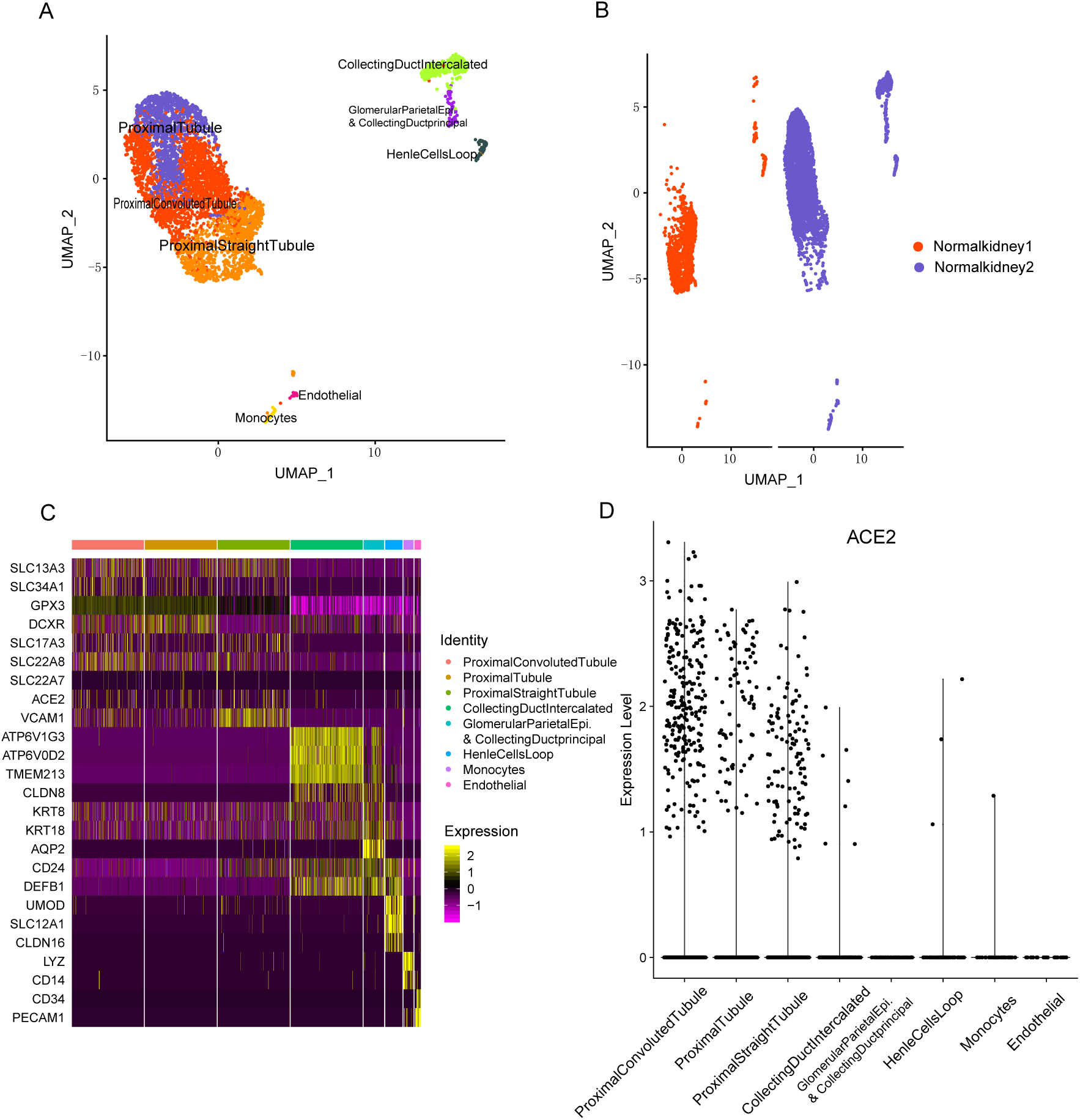
Single cell analysis of the two kidney biopsy samples. A) cell type annotations visualized by UMAP. B) The sample specific UMAP visualization of two samples. C) Heatmap of canonical markers applied for cell type assignment. D) Violin plot displaying the expression signals of kidney cell types.

As observed in the public dataset, we observed upregulated ACE2 expressions in all proximal tubule cells comparing to other cell types (Figure 2D). Quantitatively, we found about 5%-15% of both straight and convoluted proximal tubule cells expressing ACE2. Previous immunohistochemistry analysis^17^ indicated a complex spatial distribution of ACE2 protein expression, concentrated in the brush border of the proximal tubules.

### Expression pattern of ACE2 in bladder

Based on the public bladder dataset of 12 cell types (Figure 3A and B), we found low expressions of ACE2 in all epithelial cell types. The concentration of ACE2 seemed to have a decreasing trend from the out layer of the bladder epithelium (umbrella cells) to the inner layer (basal cells) with the intermediate cells in-between. Other cell types such as endothelium and immune cells are mostly negative for ACE2. The percentage of cells expressing ACE2 in umbrella cells of bladder (1.3%) is lower than that in the renal epithelium. The current reports of 2019-nCoV patients have not shown the positive detection of the virus in the urine samples. However, previous analysis of SARS patients indicated SARS virus was able to survive in urines on detectable levels ^18 19^. The detection of SARS virus in urine implied the possibility of virus releasing from infected bladder epithelial cells, in agreement with our analysis above.

**Figure 3.**
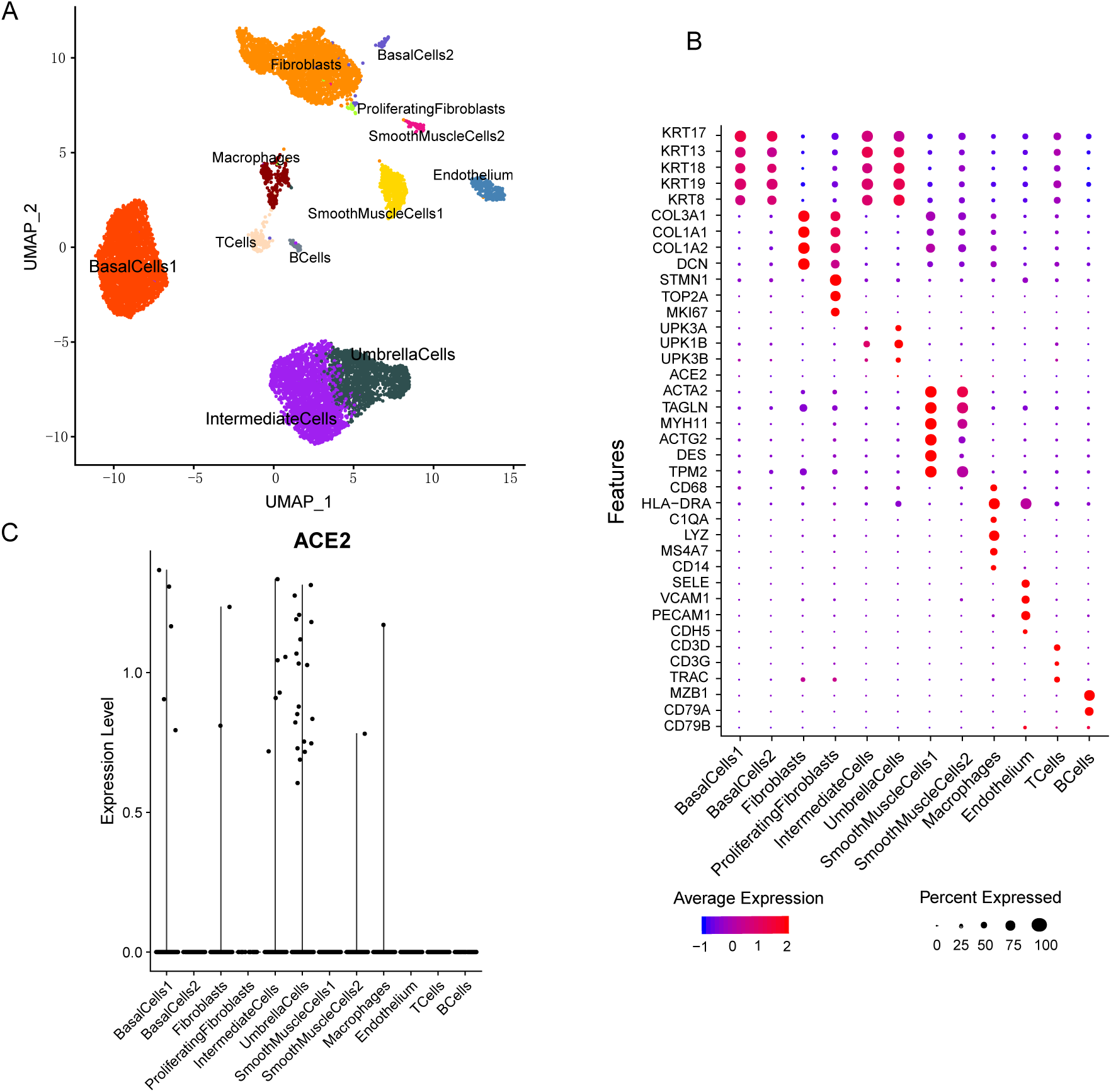
Public bladder single cell dataset. A) two-dimensional visualization of bladder cell atlas by UMAP. B) Expression pattern of canonical markers used for identifying bladder cell types. C) ACE2 expression across 12 cell subgroups in the bladder dataset.

## Discussion and Conclusion

### AKI in 2019-nCoV and SARS infection

2019-nCoV is a new infectious disease with a formidable speed of transmission and too many unknowns of pathopoiesia, which puts the world on alert since late 2019. A wide range of non-respiratory symptoms has subsequently been reported suggesting involvements of other organs during the course of the disease. In the latest analysis about the clinical features of the cases infected with 2019-nCoV, renal function damage had been confirmed^9 10^ and the continuous kidney substitution treatment (CRRT) are needed in about 9% of all 99 infected patients^9^. In addition, kidney injury and renal dysfunction had been seen in other coronaviruses associated with respiratory infections, especially two beta-coronaviruses (SARS-CoV and MERS-CoV)^20 21 22^, which are genetically similar to 2019-nCoV^4 8^. A previous study reported that 36 (6.7%) of 536 patients with SARS developed AKI and 33 of all 36 patients (91.7%) with renal dysfunction died finally^20^. Importantly, SARS -CoV fragments had been detected by PCR in urine specimens of some patients^23^. Furthermore, SARS-CoV persistent infection and replication had been observed in the kidney, particularly in tubular epithelial cells in vitro^24^, indicating that the kidney damage in patients with SARS could be owing to the direct viral infection in target cells except for the consequences of the host immune response. However, the detailed mechanism of renal involvements of 2019-nCoV and SARS-nCoV is still unclear.

### The analysis of ACE2 expression in the renal system

The etiology of AKI in SARS seems to be multifactorial and still uncertain, which was thought to have a certain correlation with the high expression of ACE2 in the kidney^24 25^. Based on our single-cell analysis in both normal kidneys and bladders, we found both detectable levels of ACE2 in kidney and bladder. Kidney PT cells have higher expression percentages than bladder epithelial cells, which may indicate that kidney is more susceptible to 2019-nCoV infection than bladder. Comparing to the expression levels of ACE2 in the digestion system ^11^ the lower expression of ACE2 in the renal and urinary system could suggest fewer patients with renal related symptoms, which positively correlates with the observations for SARS patients.

### Significance of the study

Acute kidney injury, which has been proved to be a predictor of high mortality in SARS patients^20^, may also lead to difficulty of treatment, worsening conditions and even be an negative prognostic indicator for survival for 2019-nCoV infection patients. Our study provided direct evidences for the possible infection of 2019-nCoV in the renal epithelial cells, which calls for more attention on the 2019-nCoV patients with acute kidney injury, especially during an extensive outbreak as the current status in the Hubei province. Finally, though there has been no reports of positive detection of 2019-nCoV in urine up till now, the distributions of ACE2 in the renal system and its detection in urine samples of SARS patients hinted the importance of virus-testing in urine samples of 2019-nCoV patients.

